# Monitoring of urban ecological environment including air quality using satellite imagery

**DOI:** 10.1101/2022.03.28.486114

**Authors:** Yuan Wang, Guoyin Cai, Liuzhong Yang, Ning Zhang, Mingyi Du

## Abstract

Rapid urbanisation has highlighted problems in the urban ecological environment and stimulated research on the evaluation of urban environments. In previous studies, key factors such as greenness, wetness, and temperature were extracted from satellite images to assess the urban ecological environment. Although air pollution has become increasingly serious as urbanisation proceeds, information on air pollution is not included in existing models. The Sentinel-5P satellite launched by the European Space Agency in 2017 is a reliable data source for monitoring air quality. By making full use of images from Landsat 8, Sentinel-2A, and Sentinel-5P, this work attempts to construct a new remote sensing monitoring index for urban ecology by adding air quality information to the existing remote sensing ecological index. The proposed index was tested in the Beijing metropolitan area using satellite data from 2020. The results obtained using the proposed index differ greatly in the central urban region and near large bodies of water from those obtained using the existing remote sensing monitoring model, indicating that air quality plays a significant role in evaluating the urban ecological environment. Because the model constructed in this study integrates information on vegetation, soil, humidity, heat, and air quality, it can comprehensively and objectively reflect the quality of the urban ecological environment. Consequently, the proposed remote sensing index provides a new approach to effectively monitoring the urban ecological environment.

## 1. Introduction

Many urban ecological problems have emerged with increasing urbanisation, such as the urban heat island effect, air pollution, and the degradation of water quality [1]. Although the ecological environment is the necessary material basis for human life, human activities are concentrated mainly in the urban environment. Therefore, methods of monitoring the quality of the urban ecological environment have become a focus of research on urban ecology. The information on the ecological environment is essential for urban environmental management and planning. It can provide powerful support for decision-makers and the public to understand ecological conditions and to perform urban sustainability assessments [2].

The problem of air pollution is becoming increasingly severe [3] and has attracted considerable attention in recent years. Urban ecological environment monitoring plays a fundamental role in the control of air pollution by understanding its impact on the environment [4–6]. The main gaseous pollutants affecting urban air quality include trace gases such as nitrogen dioxide, sulfur dioxide, carbon monoxide, and ozone. Among them, nitrogen dioxide is a very serious air pollutant, and it is also among the main pollutants during heavy air pollution weather in China [7].

With the development of satellite remote sensing, satellite image data have played an important role in urban ecological environment monitoring because of their high spatial resolution, wide coverage, and fast acquisition [8–10]. On October 13, 2017, the European Space Agency successfully launched the Sentinel-5P satellite for air quality monitoring [11,12]. Its Tropospheric Monitoring Instrument can provide global daily coverage [11] and collect near-real-time air quality data [13]. Sentinel-5P is the first mission dedicated to monitoring the atmosphere, and it has proved to be effective in monitoring air pollutants with support from Sentinel-2 and other supplementary data (https://www.esa.int/Applications/Observing_the_Earth/Copernicus/Sentinel-5P/Mapping_methane_emissions_on_a_global_scale).

Many researchers have measure ecological parameters for various purposes, such as the evaluation of ecological vulnerability [14–16], ecological safety evaluation [17–19], efficiency evaluation [20–22], and the evaluation of the ecological environment carrying capacity [23–25]. Xu [26] proposed the remote sensing ecological index (RSEI), which can be used for remote sensing monitoring and the comprehensive evaluation of a regional ecological environment. The RSEI has many advantages, such as capacities for visualisation, spatiotemporal analysis, and quantitative description. By combining the RSEI and ecological sensitivity analysis, Yang et al. [27] proposed a Markovian model to delineate the urban growth boundary in the ecologically fragile areas of the Upper Yellow River of China. Firozjaei et al. [28] proposed an additional analytical framework for assessing the surface ecological status in urban environments by considering five biophysical characteristics: surface greenness, dryness, moisture, heat, and imperviousness. Because the effect of air quality on the ecological environment is increasing, this work considers the air pollution index (API) and develops an air-quality-included RSEI (AQRSEI) based on the RSEI. The proposed index is calculated and analysed for the administrative region of Beijing. The remainder of this paper is organised as follows. Section 2 introduces the study area, and Section 3 describes the data sources and method. Results and analyses are presented in Section 4, and the results are discussed in Section 5. Section 6 concludes the paper.

## 2. Study area

As the capital city of China, Beijing is a central area integrating national politics, culture, international exchange, and scientific and technological innovation [29]. Beijing consists of 16 municipal districts covering an area of 115.7°–117.4°E and 39.4° –41.6°N. The centre is located at 116°20′E and 39°56′N, and the total area is 16,410.54 km2. The terrain of Beijing is high in the northwest and low in the southeast, with an average altitude of 43.5 m. The climate is a warm temperate semi-humid and semi-arid monsoon climate, with high temperatures and rain in summer, cold dry weather in winter, and short transitional periods in spring and autumn.

Because most human activities are concentrated in urban areas, the ecological environment status is vitally important for people living in urban region [30]. As the capital of China and a world megacity, Beijing has gradually faced difficulties caused by urban problems such as traffic congestion, shortage of water resources, and air pollution [31]. In this study, Beijing is taken as the research area, and its urban ecological environment is quantitatively analysed. The results can provide useful information for decision-making on urban environmental protection by local governments and contribute to the sustainable development of Beijing [32]. Fig 1 shows a Landsat-8 remote sensing image of the study area; the red line is the administrative boundary of Beijing.

**Fig 1.**
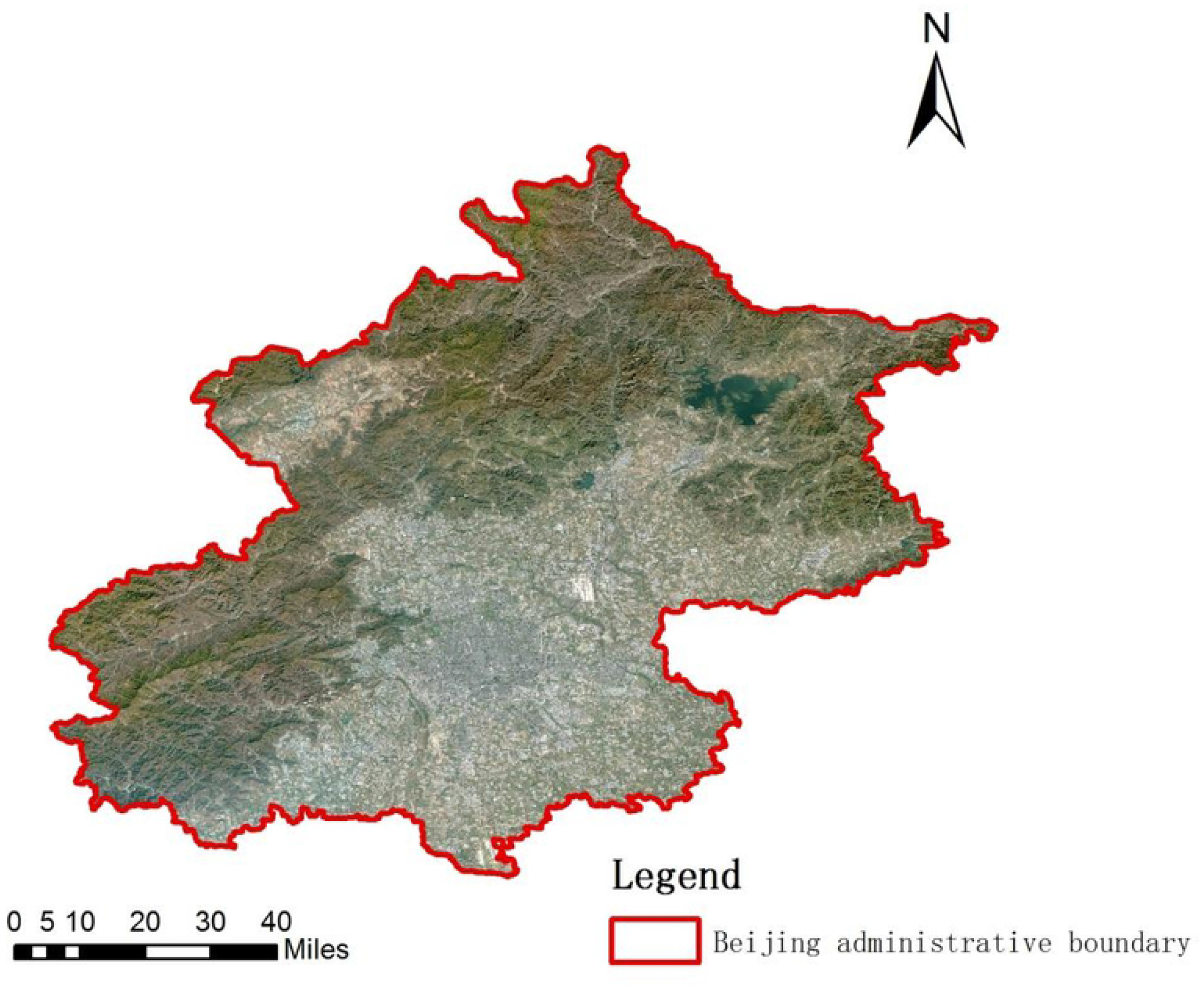
False-colour composite map of the study area from Landsat-8 images of Beijing, 2020.

## 3. Materials and methods

### 3.1 Data Sources and Processing

This study evaluates the ecological environment of Beijing using five indices: green-ness, dryness, wetness, temperature, and air pollution. Remote sensing images from Sen-tinel-2A, Landsat-8, and Sentinel-5P in 2020 were used as data sources, and the global urban boundary data released by Tsinghua University were used as the urban boundary of Beijing. Sentinel-2A images were used to calculate the greenness, dryness, and wetness, and Landsat-8 thermal infrared sensor (TIRS) data were employed to compute the temperature. Air pollution information was extracted from Sentinel-5P data. In addition to spatial data, other materials such as the local ecological environment quality report “ Communique on the State of Beijing’s Ecological Environment in 2020” and national environmental quality standard “Environmental Quality Standards for Surface Water” were employed as supplementary data. Table 1 describes the satellite data.

**Table 1a.** Band characteristics of Sentinel-2A and Landsat-8used in this work.

| Satellite | Band Number | Pixel Size | Wavelength | Description |
| --- | --- | --- | --- | --- |
| Sentinel-2A | B1 | 60 | 0.443 | Coastal Aerosol |
|  | B2 | 10 | 0.490 | Blue |
|  | B3 | 10 | 0.560 | Green |
|  | B4 | 10 | 0.665 | Red |
|  | B5 | 20 | 0.705 | Vegetation Red Edge |
|  | B6 | 20 | 0.740 | Vegetation Red Edge |
|  | B7 | 20 | 0.783 | Vegetation Red Edge |
|  | B8 | 10 | 0.842 | NIR |
|  | B8A | 20 | 0.865 | Vegetation Red Edge |
|  | B9 | 60 | 0.945 | Water Vapor |
|  | B10 | 60 | 1.375 | SWIR–Cirrus |
|  | B11 | 20 | 1.610 | SWIR 1 |
|  | B12 | 20 | 2.190 | SWIR 2 |
| Landsat-8 | B10 | 100 | 10.60–11.19 | TIRS 1 |
|  | B11 | 100 | 11.50–12.51 | TIRS 2 |

**Table 1b.** Characteristics of Sentinel-5P bands used in this work.

|  | Spectral Band | Spectral Range (nm) | Spectral Resolution (nm) |
| --- | --- | --- | --- |
| Sentinel-5P | UV 1 | 270–300 | 0.5 |
|  | UV 2 | 300–320 | 0.5 |
|  | UV–VIS | 310–405 | 0.55 |
|  | VIS | 405–500 | 0.55 |
|  | NIR 1 | 675–725 | 0.5 |
|  | NIR 2 | 725–775 | 0.5 |
|  | SWIR | 2305–2385 | 0.25 |

Because the greenness index is closely related to vegetation and the dryness and wetness indices are closely related to soil, buildings, and water bodies, Sentinel-2A images can be used not only to monitor the growth of vegetation, but also to extract information about soil and water bodies [33]. Landsat-8 is equipped with an operational land imager and TIRS; it provides global remote sensing data from April 2013 to the present with a spatial resolution of 30 m. The T1 image of Landsat-8 released by the U.S. Geological Survey is used in this work. The B10 and B11 bands are brightness temperature bands with a resolution of 100 m (resampled to 30 m); they can be used to retrieve the temperature index. Sentinel-5P can accurately monitor the concentrations of trace gases such as ozone, carbon monoxide, nitrogen dioxide, sulfur dioxide, and methane in the atmosphere [34–36], which can reflect the air pollution in the study area.

To acquire cloud-free remote sensing images of the study area over time and space, the image data must be preprocessed. The Q60 band of the Sentinel-2 satellite is described as a cloud mask and is used for cloud removal. Landsat surface reflectance images can be processed using the pixel_qa band for cloud removal. However, Landsat top-of-atmosphere data have no information in the pixel_qa band; therefore, the statistical mono-window (SMW) algorithm [37] was used for cloud removal in this dataset.

Google Earth Engine (GEE) is an online visual cloud computing platform. It provides massive global-scale satellite data, and it has sufficient computing power to provide powerful data computing, analysis, storage, and management capabilities [38]. There is no need to download the image data, which can be processed and analysed online, and existing code data can be saved online. The GEE platform was selected for data processing, model construction, index calculation, and results analysis.

### 3.2 Method

#### 3.2.1 Methods of calculating the indices

The RSEI is an ecological evaluation model that couples four urban ecological elements: greenness, dryness, wetness, and temperature [39]. They are identified as the vegetation index, soil index, humidity component of the tasseled cap transformation, and sur-face temperature, respectively, and can be calculated from remote sensing data. Principal component analysis (PCA) is used to integrate each index, and the results can be used to quantify the ecological environment quality.

In addition to these four indices, the proposed AQRSEI considers information on air quality. All of these indicators are closely related to urban ecological quality. Because the indicators are obtained from remote sensing images, they can be used to accurately and objectively describe the ecological environment in the study area [39].

##### 1. Normalised difference vegetation index

The normalised difference vegetation index (NDVI) reflects the growth state and coverage of vegetation. In this study, it was used to characterise the greenness index. The NDVI generated by Sentinel-2A accurately reflects vegetation greenness, photosynthesis intensity, and vegetation metabolism intensity [40,41] and can be used to monitor and analyse vegetation and crops in cities. It is calculated as follows:

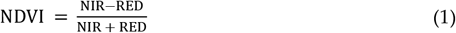

where NIR and RED represent the reflectance in the near-infrared and red bands, respectively.

##### 2. Normalised difference building-soil index

The normalised difference building-soil index (NDBSI), which is generated using a combination of the bare soil index (SI) and the index-based built-up index (IBI), was used to represent the degree of soil desiccation in the study area [42]. The main factors affecting urban dryness are soil and buildings in the city. Therefore, it is reasonable to use the NDBSI for calculation. It is calculated as follows using the average values of the SI and the IBI.

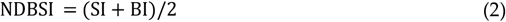

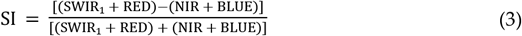

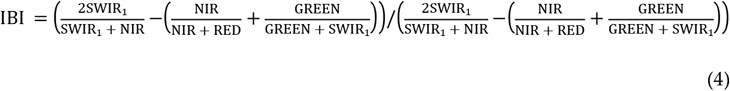

where SWIR1 and BLUE represent the reflectance in the shortwave infrared-1 and blue bands, respectively.

##### 3. Modified normalised difference water index

The study area includes four large reservoirs (Mi-Yun, Guanting, Huairou, and Haizi reservoirs), five major river systems (the Yongding, Chaobai, Beiyun, Juma, and Jiyun rivers), and other rivers with abundant groundwater resources. The modified normalised difference water index (MNDWI) was used as the wetness index in this work. It is quicker to apply than the normalised difference water index (NDWI) and is more suitable for water body information extraction in cities [43]. It is calculated as follows:

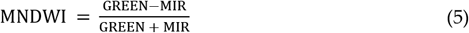

where GREEN and MIR represent the reflectance in the green and mid-infrared bands, respectively.

##### 4. Land surface temperature

Land surface temperature (LST) is used to measure urban regional temperature. The SMW algorithm and Landsat-8 satellite image data were used to calculate the heat component. The algorithm expresses the empirical relationship between the atmospheric ap-parent brightness temperature and the surface temperature obtained from a single thermal infrared band through a simple linear relationship. LST is calculated as follows:

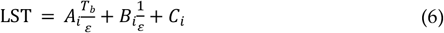

where Tb is the brightness temperature in the thermal infrared channel, ε is the emissivity of that channel, and Ai, Bi, and Ci are algorithm coefficients.

##### 5. Air pollution index

The Sentinel-5P satellite was designed to perform near-real-time atmospheric measurement tasks, and its data can be used to monitor air quality, ultraviolet radiation, gas concentrations, and air pollutants. The main urban air pollutants are carbon monoxide (CO), nitrogen oxides (NOx), ozone (O3), sulfur dioxide (SO2), hydrocarbons, and particulate matter (PM2.5 and PM10). The main source of nitrogen dioxide (NO2) is automobile exhaust emissions, industrial emissions, and biological combustion. Its concentration in cities is relatively high [44]. As a result, the API considers four pollutant gases: NO2, SO2, O3, and CO. Following one study [45], weights are assigned according to the degree of harm caused by each pollutant gas: NO2 accounts for 70%, SO2 accounts for 15%, O3 accounts for 10%, and CO accounts for 5%. The four gases are integrated into one air pollution component, as follows:

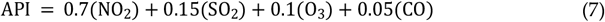

#### 3.2.2 Principal components analysis

PCA is a data dimensionality reduction algorithm. The calculation method is simple and easy to implement. It integrates various indicators into a few new indicators. These indices are considered comprehensively by preserving as much of the original information as possible. Because the dimensions of the five indicators are not uniform, the indicators must be normalised before PCA is performed.

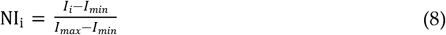

where represents the value of an index after normalisation, and and represent the maximum and minimum values of an index, respectively.

After each index is normalised, PCA can be applied to the five indices together. Ac-cording to a previous study [39], the first component extracted in a PCA (PC1) represents most of the information in the input data. PC1 is inversely related to the ecological environment quality; by subtracting PC1 from 1, a result with a direct relationship with the ecological environment quality is obtained, and thus the value 1 – PC1 can be regarded as the initial AQRSEI, AQRSEI0; higher values of AQRSEI0 indicate better ecological environment quality. AQRSEI0 can be expressed as

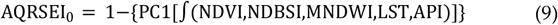

AQRSEI0 is normalised to obtain the AQRSEI as follows:

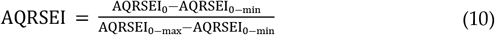

The AQRSEI is divided into five levels. Values between 0 and 0.2 indicate poor ecological quality, and values between 0.2 and 0.4 indicate relatively poor quality. Moderate quality is indicated by values between 0.4 and 0.6. The ranges of 0.6– 0.8 and 0.8–1 indicate relatively high and high ecological quality, respectively.

## 4. Results and analysis

### 4.1 Result

On the GEE platform, using the calculation methods described above, the NDVI, NDBSI, and MNDWI were calculated using Sentinel-2A data for Beijing in 2020, the LST was calculated using Landsat-8 data, and the API was calculated using Sentinel-5P data. Fig 2 shows the normalised greenness (Fig 2-a), normalised dryness (Fig 2-b), normalised wetness (Fig 2-c), normalised temperature (Fig 2-d), and normalised air pollution (Fig 2-e). According to the Fig 2-a, the overall level is high in most of the study area but significantly lower in the central area; the reason is the characteristics of the subsurfaces in the study area, where mountains surround the central urban region.

**Fig 2.**
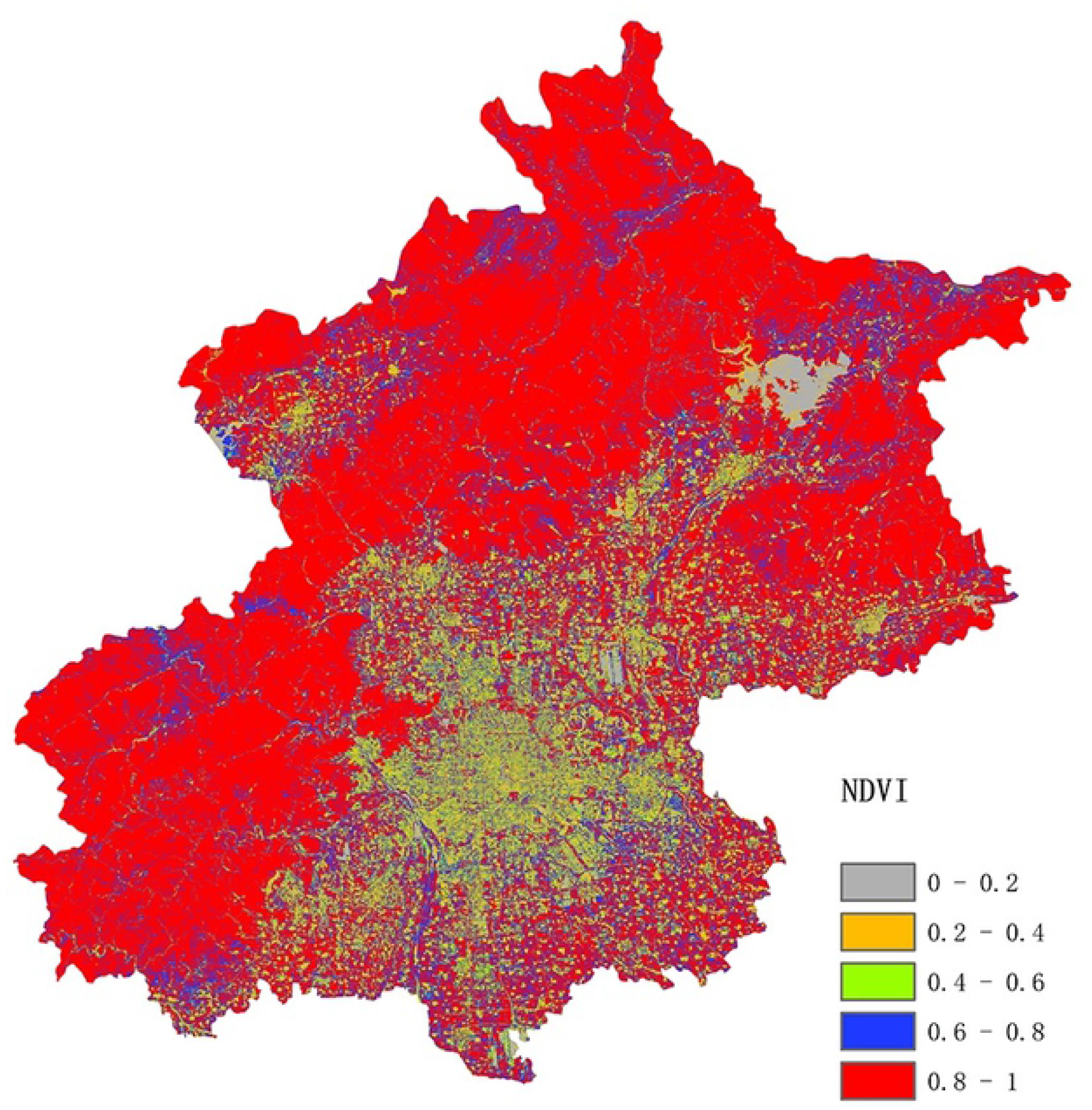
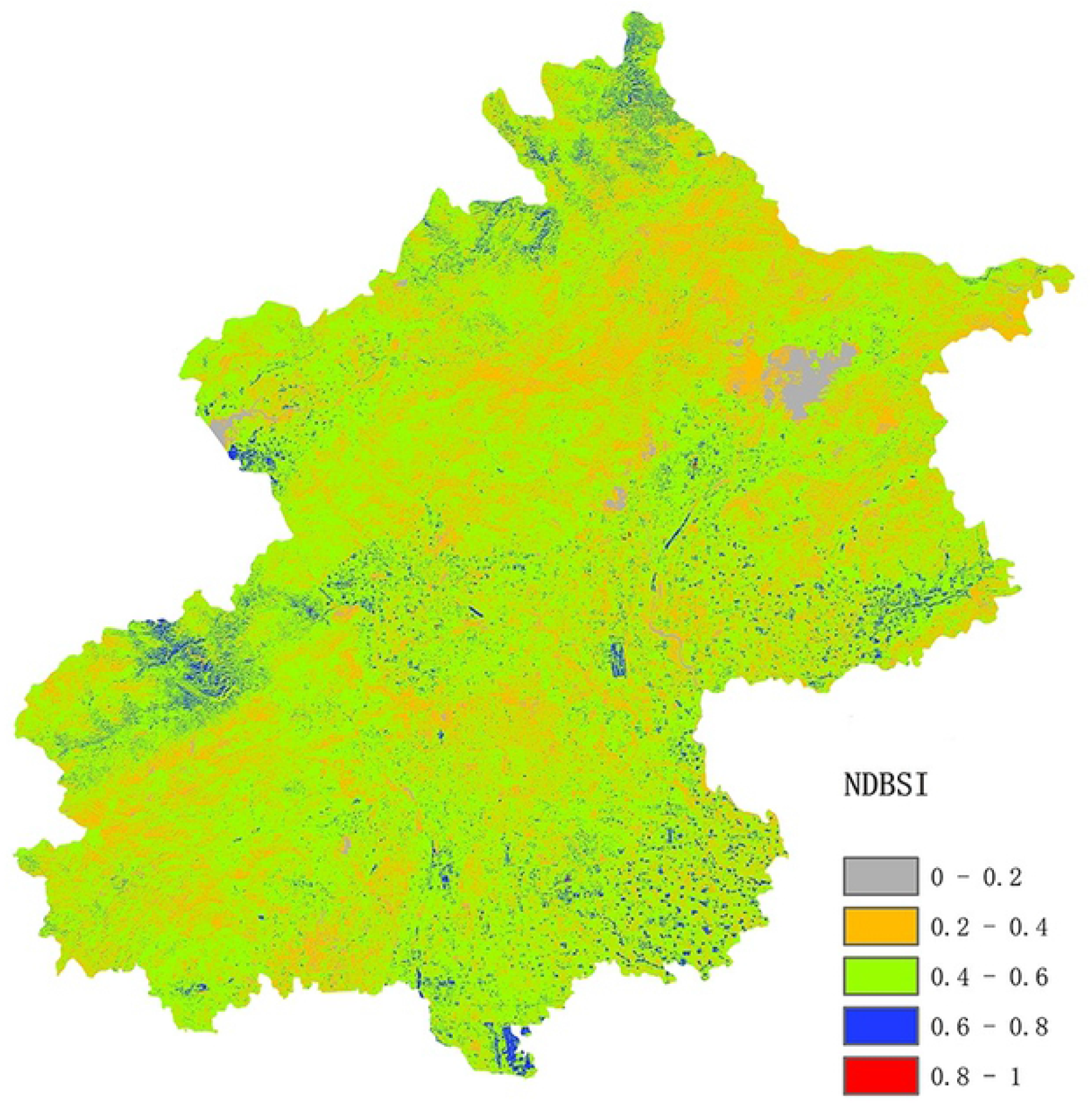
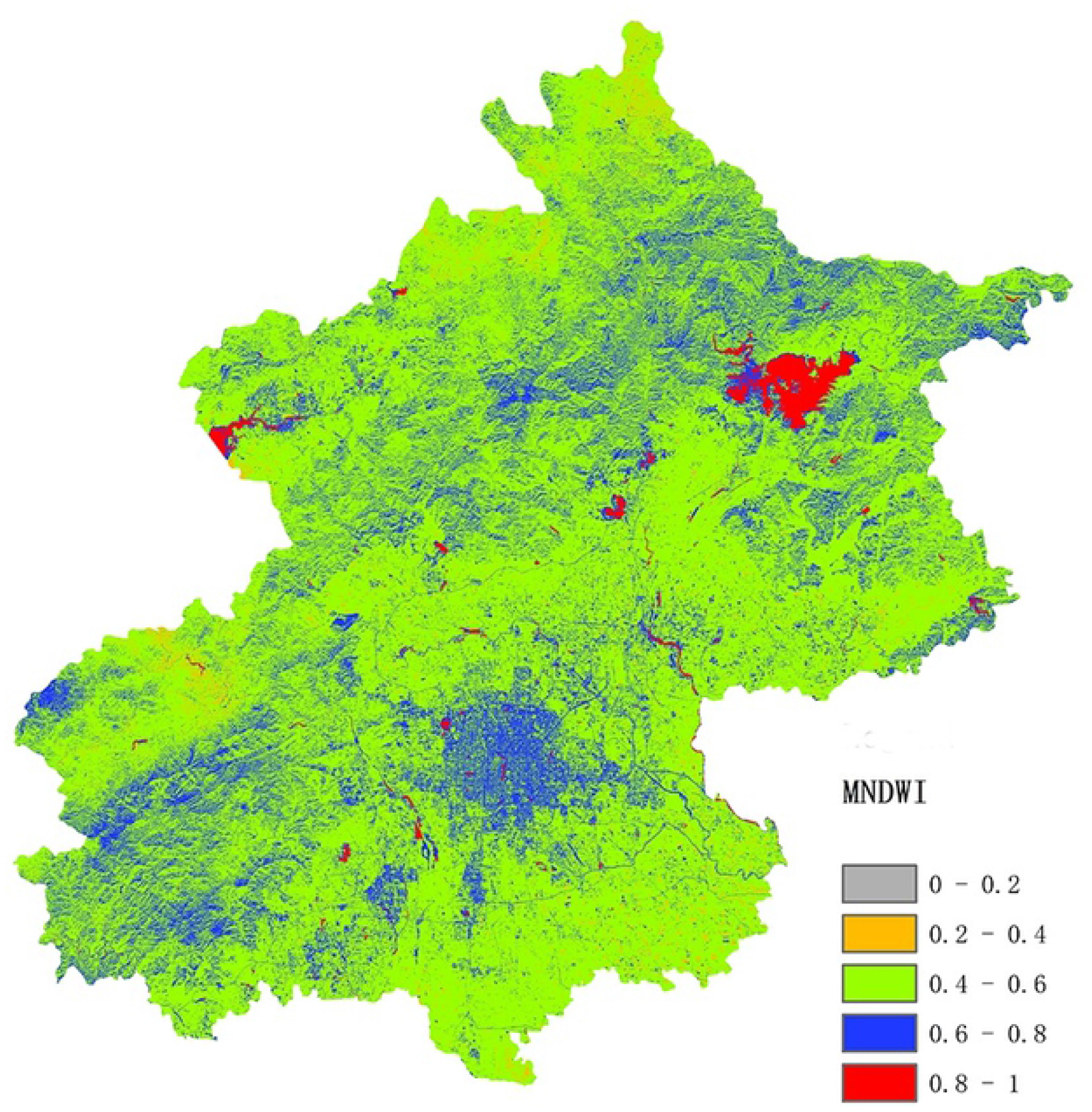
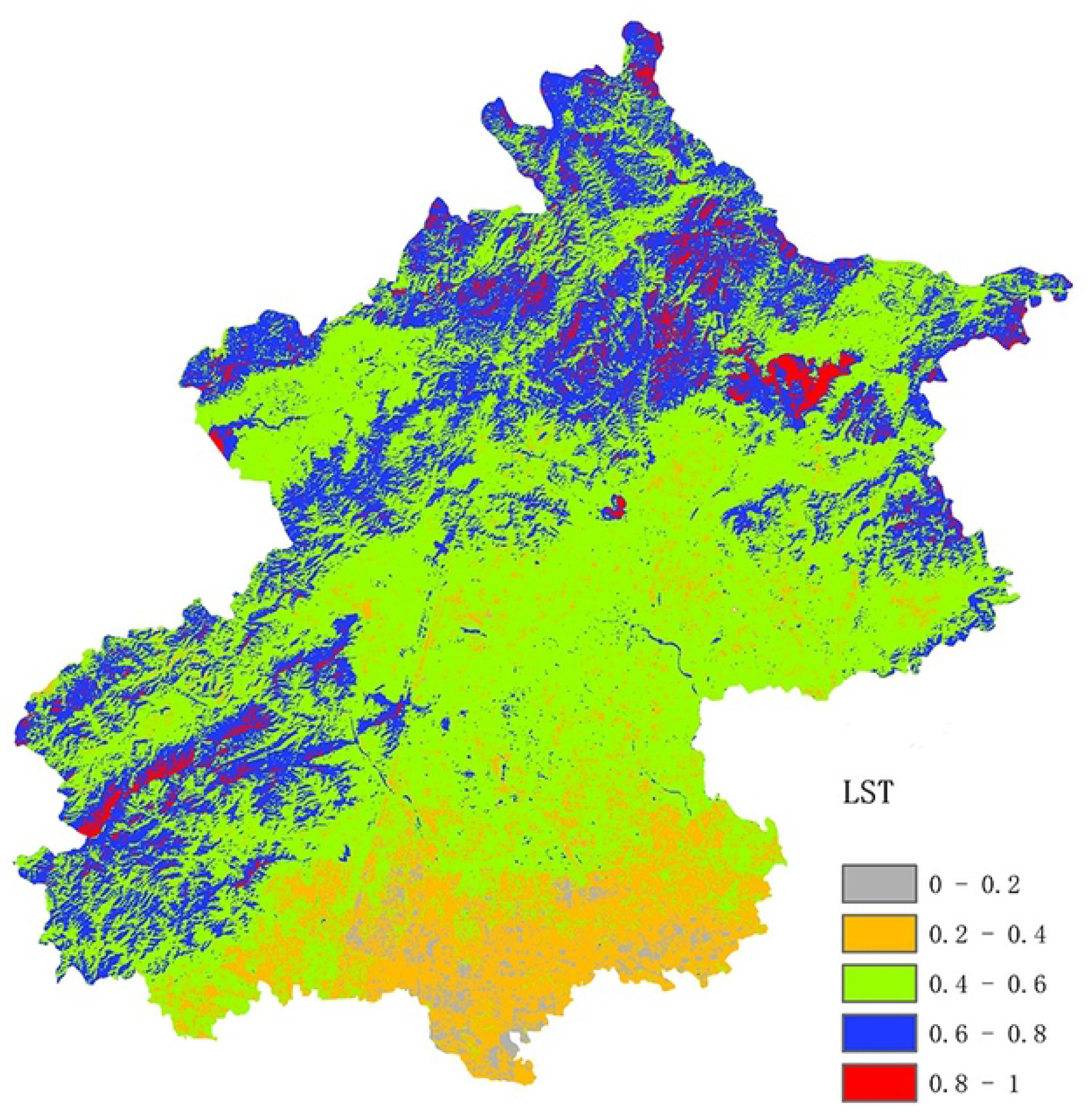
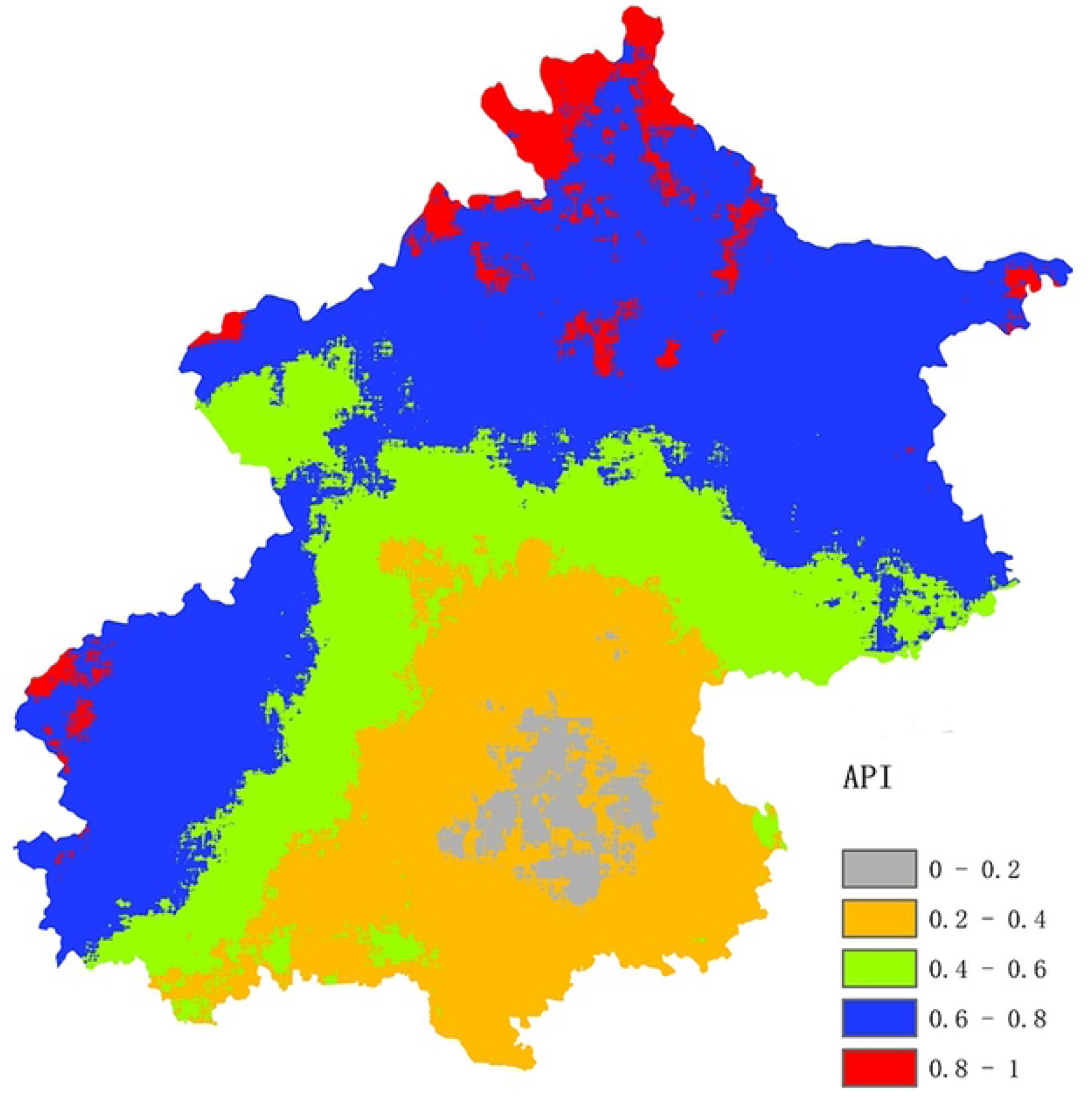
Maps of normalized (a) greenness, (b) dryness, (c) wetness, (d) temperature, and (e) air pollution in the study area. **a** Normalised greenness map, **b** Normalised dryness map, **c** Normalised wetness map, **d** Normalised temperature map, **e** Normalised air pollution map

Fig 2-b shows that overall the study area is moderately dry, and the distribution is relatively uniform, with no regular difference; this result is related to the distribution of bare soil and impervious surfaces. Fig 2-c shows that areas near reservoirs have the highest wetness; the central urban area and most parts of the mountainous areas have good wetness, and the wetness in the study area is good overall. The Fig 2-d shows a trend of higher LST in the southern part of the study area. There is a clear LST difference between the north and south because there are factories in the southern part of Beijing. Unlike the other indicators, the Fig 2-e shows a roughly circular structure. The central urban area is severely polluted, and rings of increasingly higher air quality appear outside of the central area; this pattern is closely related to the land use/cover in Beijing, where the central areas are built up, and the surrounding regions consist of farm land and forests.

The five indices (greenness, dryness, wetness, temperature, and air pollution) were normalised before PCA was performed. The PCA results are shown in Table 2. There are five principal components, PC1–PC5. The eigenvalue, eigenvalue contribution rate, variance, standard deviation, and mean value of the AQRSEI are also given.

**Table 2.** PCA results for AQRSEI in 2020.

| **Year** | **2020** |  |  |  |  |
| --- | --- | --- | --- | --- | --- |
| **Index** | **PC1** | **PC2** | **PC3** | **PC4** | **PC5** |
| **Greenness** | 0.9367 | -0.3344 | -0.0990 | -0.0383 | -0.0072 |
| **Dryness** | -0.3201 | 0.1401 | -0.8796 | 0.2906 | -0.1658 |
| **Wetness** | 0.1108 | -0.0388 | 0.9538 | -0.1880 | -0.2298 |
| **Temperature** | -0.7890 | -0.0680 | -0.3910 | -0.4665 | -0.0155 |
| **Air Pollution Index** | -0.8280 | -0.5531 | 0.0537 | 0.0719 | 0.0013 |
| **Eigenvalue** | 0.0849 | 0.0181 | 0.0135 | 0.0041 | 0.0005 |
| **Eigenvalue Contribution Rate** | 70.11% | 14.95% | 11.15% | 3.39% | 0.41% |
| **Variance** | 0.0565 | 0.0074 | 0.0055 | 0.0138 | 0.0378 |
| **Standard Deviation** | 0.2378 | 0.0860 | 0.0742 | 0.1175 | 0.1944 |
| **Mean Value of AQRSEI** | 0.6793 |  |  |  |  |

The contribution rate of the PC1 eigenvalue exceeds 70%, indicating that PC1 represents most of the eco-environmental information in the five indicators. Greenness and wetness show positive values of PC1, indicating that they have a positive impact on the urban environment, whereas the values for dryness, temperature, and air pollution are negative, indicating that they negatively affect the urban environment. According to the absolute value of PC1, these five indicators affect the ecological environment in the following order: greenness > air pollution > temperature > dryness > wetness. This result indicates that air pollution plays a key role in urban environmental assessment and demonstrates the effectiveness of the API. The average value of the index for remote sensing monitoring of the ecological environment can indicate the average ecological environment quality of a region [46]. The average AQRSEI obtained in this study is 0.6793, indicating that the ecological quality in this area is at a relatively high level.

### 4.2 Comparison with RSEI

The PCA results for the existing RSEI were calculated for comparison and are presented in Table 3.

**Table 3.** PCA results for RSEI in 2020.

| **Year** | **2020** |  |  |  |
| --- | --- | --- | --- | --- |
| **Index** | **PC1** | **PC2** | **PC3** | **PC4** |
| **Greenness** | 0.9891 | -0.1300 | -0.0748 | 0.0074 |
| **Dryness** | -0.3605 | -0.8535 | 0.3438 | 0.1666 |
| **Wetness** | 0.1354 | 0.9383 | -0.2370 | 0.2292 |
| **Temperature** | -0.7594 | -0.4094 | -0.5036 | 0.0135 |
| **Eigenvalue** | 0.0643 | 0.0135 | 0.0050 | 0.0005 |
| **Eigenvalue**<br>**Contribution Rate** | 77.19% | 16.20% | 6.00% | 0.60% |
| **Variance** | 0.0565 | 0.0074 | 0.0055 | 0.0138 |
| **Standard Deviation** | 0.2378 | 0.0860 | 0.0742 | 0.1175 |
| **Mean Value of RSEI** | 0.6983 |  |  |  |

According to Table 3, greenness and temperature strongly affect the environment. The contribution rate of PC1 for the RSEI is 77.19% (7.08 points higher than that for the AQRSEI), and the mean RSEI is 0.6983 (0.019 higher than that of the AQRSEI). The variance and standard deviation of PC1 are consistent with the results for the AQRSEI. The eigenvalue contribution rate of PC1 is also significantly higher than that of the other indicators, indicating that the results for the AQRSEI and RSEI are similar overall, and the evaluation indicates the same eco-environmental quality level. However, in the proposed index, the API is in second place, after greenness, in terms of their individual impact on the urban environment, indicating that the proposed air quality factor has a greater impact on the environment and ultimately decreases the average value of the overall ecological index.

Fig 3 shows the PCA results of the RSEI and AQRSEI of Beijing in 2020. The results are similar overall, although differences appear in local areas, for example, the area inside the green box in the upper right corner of the Fig. The Mi-Yun Reservoir is located in this area. The water quality in this reservoir meets the class II standard for the environmental quality of surface water in China. In the RSEI diagram, the overall level of the reservoir is between 0.2 and 0.4, indicating poor quality. In the AQRSEI diagram, the overall level of the reservoir is between 0.4 and 0.6, indicating moderate quality. In light of the quality standard of the reservoir given above, the AQRSEI result is closer to the true state of the ecological environment. In addition, the yellow box in the upper left corner of the Fig contains most of Yanqing District, which has continuously promoted the construction of ecological infrastructure and improved the quality of urban construction in the suburbs of Beijing in recent years. This part of the RSEI map does not show any notable features; by contrast, in the AQRSEI result, the central area is in the range 0.6–0.8 (blue), indicating relatively high ecological environment quality. Furthermore, in the AQRSEI assessment, the mountainous area is in the range 0.8–1 (red) overall, indicating high ecological environment quality. In the outer suburbs, the urban counties are in the range 0.6–0.8 (blue) overall, indicating relatively high quality, whereas the urban area is in the range 0.4–0.6 (green) overall, indicating moderate quality. In the central part of the central urban area, the environmental quality is poor 0.2–0.4 (yellow), and the overall pattern shows a clear circular distribution. By contrast, in the RSEI map, the urban area shows notable image noise with no significant regular distribution. This result shows that the AQRSEI evaluation provides better results than the RSEI evaluation. Additionally, the map obtained using the AQRSEI is visually superior that obtained using the RSEI, which shows an obvious salt-and-pepper effect, especially in the central urban areas. The improved map quality is attributed to the calculation of the API. Therefore, the AQRSEI can measure the urban ecological environment quality from a broader perspective.

**Figure 3.**
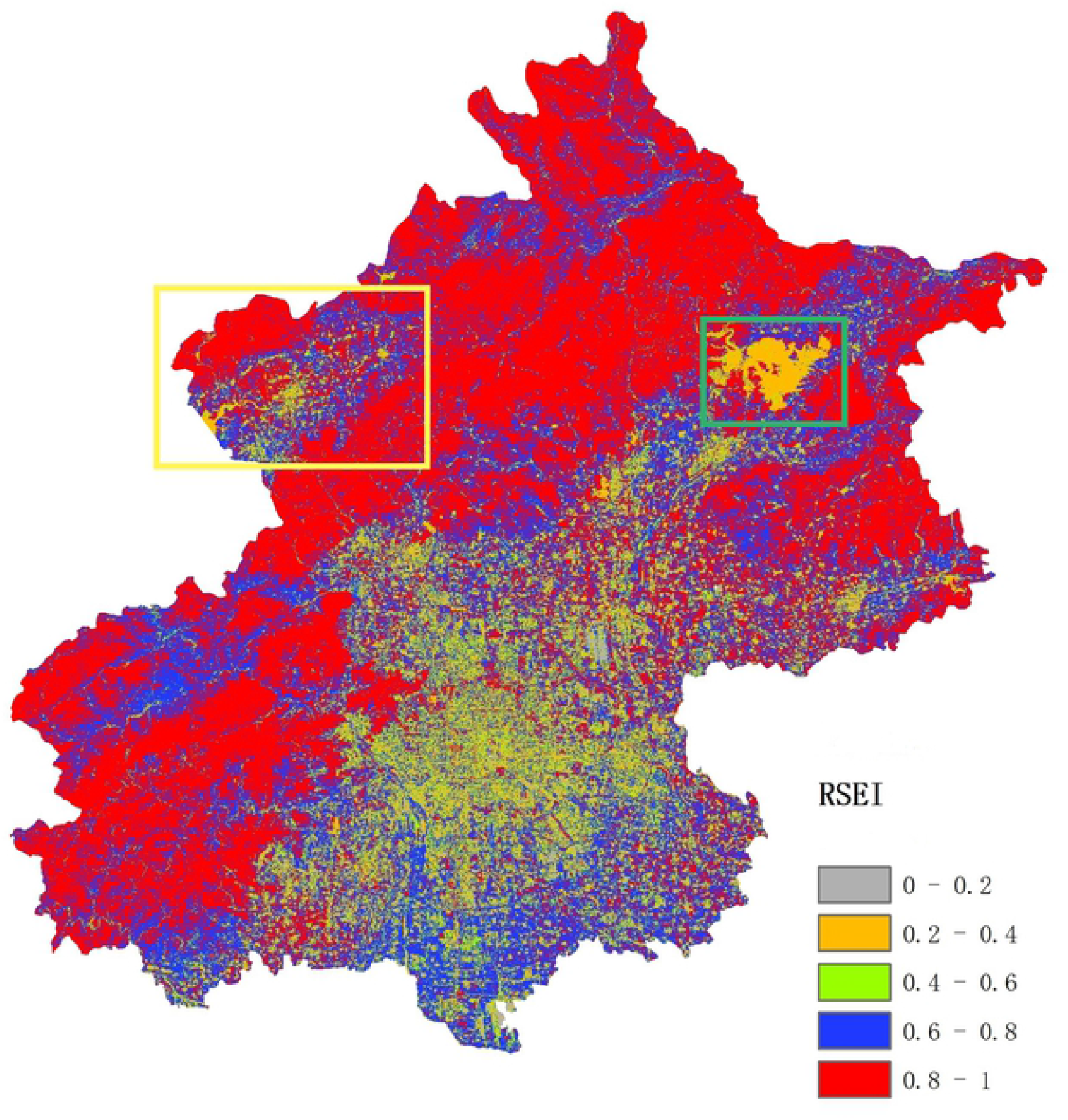
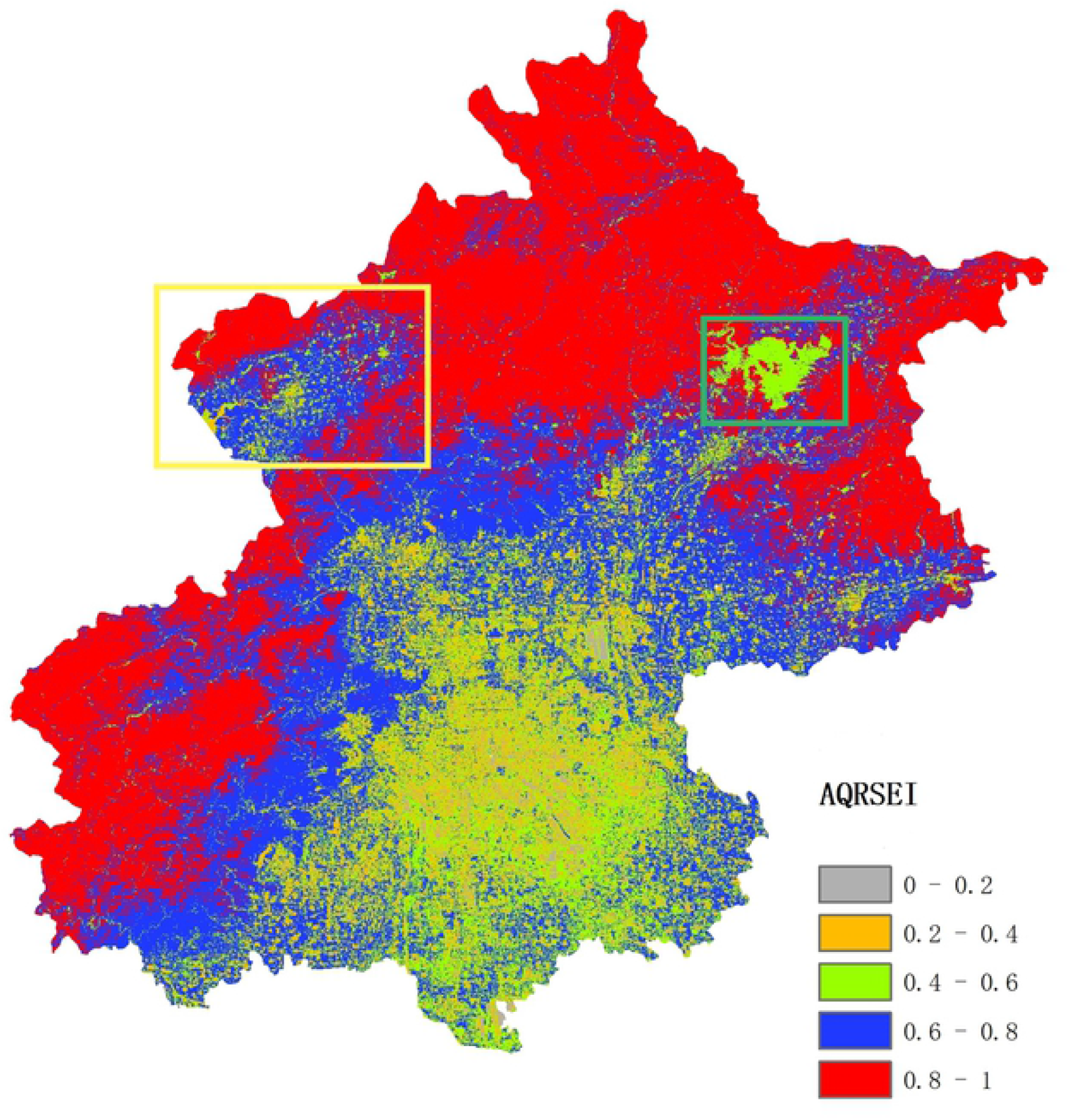
Results of PCA for (a) RSEI and (b) AQRSEI.

## 5. Discussion

Because most human activities are concentrated in urban areas and air pollution increases with rapid urbanisation, information on air pollution plays a key role in the detection, monitoring, and assessment of the urban ecological environment. Most earlier satellite-based ecological environment models were based on indicators such as greenness, wetness, dryness, and temperature, whereas the proposed index includes the factor of air pollution. Compared with those of the RSEI proposed by Xu [26], the results show that the proposed AQRSEI can provide a better indication of the true situation than the RSEI. In Beijing, the values of the remote sensing ecological index for mountainous areas, suburbs, and urban areas differ, indicating an ecological environment quality ranking of mountainous areas > suburbs > urban areas. This result is consistent with the actual situation of the ecological environment in the study site. The PCA results show that the greenness and wetness indices are positive, indicating that these factors have a positive effect on the environment. By contrast, the dryness, temperature and air pollution indices are negative, indicating that they have a negative effect on the environment. These results are consistent with those of the RSEI. In addition, the results for the PC1 component obtained by PCA show that the proposed AQRSEI outperforms the RSEI in typical areas and in the region as a whole, indicating that the model constructed in this study is reliable.

The study area in this work covers the entire administrative region of Beijing. It is a local-scale case study focused on the RSEI model; therefore, accessible satellite data from Sentinel-2A, Landsat-8, and Sentinel-5P for 2020 were employed. GEE was used to acquire satellite images, develop the model, and calculate the indices. This method uses accessible time-series satellite data, and the algorithms are applied quickly over large spatial extents. A reasonable multi-source data fusion evaluation was performed using remote sensing image data from different satellites, thus combining the advantages of each satellite. Moreover, many aspects of the ecological environment of the study area were objectively monitored and evaluated. However, it is time-consuming to search for cloud-free satellite data over large areas. Although the GEE platform includes cloud removal algorithms and mosaic operations, the processing of the images will negatively affect the results of the index calculation.

The addition of the API will affect the overall results because its impact is negative and the value is relatively large, indicating that it has a large negative impact on the urban environment and thus affects the overall AQRSEI mean value. Because the establishment of the API depends on the weights assigned in a single measurement index according to the degree of harm of pollutant gases, it may cause human factor interference and consequently influence the results. This problem is also a key point that requires detailed in-depth analysis in future work.

There are two possible approaches to further research. The first approach is to strengthen the API to make it more truly and accurately reflect air quality, for instance, by adding information on particulate matter (PM10, PM2.5), which is important in air quality studies. The second is to introduce other indicators such as the water pollution index and percentage of impervious surfaces to monitor the ecological environment and develop a satellite-based model for measuring environmental parameters more comprehensively and objectively.

## 6. Conclusions

Owing to the rapid pace of economic growth and urbanisation, improved living standards have attracted public attention to environmental quality. It is particularly important to monitor and evaluate the living environment. This study used the GEE cloud computing platform and added an air pollution component to the existing RSEI to analyse the ecological environment of Beijing in 2020. The main conclusions are as follows.

The proposed index includes five indicators: greenness, dryness, wetness, surface temperature, and API. The eigenvalue contribution rate of PC1 for the proposed AQRSEI model is greater than 70%, which indicates that the first component obtained by PCA represents most of the information of each index. According to the values of PC1, the strength of the effects of the five indicators on the environment follows the order greenness > air pollution > temperature > dryness > wetness. This result indicates that the air quality index can play a key role in the assessment of the ecological environment.

The results from both the RSEI and the AQRSEI indicate that the ecological status of Beijing in 2020 was relatively high. However, ecological environment maps showed that the AQRSEI gives more reasonable results for water bodies and the central urban region of the study area than the RSEI. Additionally, the map based on the AQRSEI exhibits better quality with less salt-and-pepper effect than that based on the RSEI. Information on the urban ecological environment can be obtained quickly and efficiently using multi-source remote sensing images by means of the GEE cloud computing platform. In the future, more relevant factors for ecological environment assessment should be considered to develop a more comprehensive model and effectively assess the urban ecological environment.

Air quality was measured as gaseous pollution in this work; however, PM has caused widespread concern because of its critical negative impact on air quality, human health, and the natural environment in recent years. Therefore, information on PM pollution from satellite images will be considered in further studies. In addition, other factors such as a water pollution index and the percentage of impervious surface will be included to develop a satellite-based RSEI model for more comprehensive and objective measurement of the ecological environment.

## Acknowledgments

This work is financially supported by National Natural Science Foundation (NSFC) (Key Project #41390650), and National Key Research and Development Program of China (No. 2018YFC0706003).

